# PREDetector 2.0: Online and Enhanced Version of the Prokaryotic Regulatory Elements Detector Tool

**DOI:** 10.1101/084780

**Authors:** Pierre Tocquin, Aymeric Naômé, Samuel Jourdan, Sinaeda Anderssen, Samuel Hiard, Gilles P. van Wezel, Marc Hanikenne, Denis Baurain, Sébastien Rigali

**Affiliations:** InBioS – PhytoSYSTEMS, University of Liège, Liège, 4000, Belgium; Hedera-22 SCRL, Esneux, 4130, Belgium; InBioS – Center for Protein Engineering, University of Liège, Liège, 4000, Belgium; Systems and Modeling - Montefiore Institute, University of Liège, Liège, 4000, Belgium; Microbial Biotechnology - Institute of Biology, Leiden University, Leiden, 2333 BE, The Netherlands

## Abstract

In the era that huge numbers of microbial genomes are being released in the databases, it becomes increasingly important to rapidly mine genes as well as predict the regulatory networks that control their expression. To this end, we have developed an improved and online version of the PREDetector software aimed at identifying putative transcription factor-binding sites (TFBS) in bacterial genomes. The original philosophy of PREDetector 1.0 is maintained, *i.e.* to allow users to freely fix the DNA-motif screening parameters, and to provide a statistical means to estimate the reliability of the prediction output. This new version offers an interactive table as well as graphics to dynamically alter the main screening parameters with automatic update of the list of identified putative TFBS. PREDetector 2.0 also has the following additional options: (i) access to genome sequences from different databases, (ii) access to weight matrices from public repositories, (iii) visualization of the predicted hits in their genomic context, (iv) grouping of hits identified in the same upstream region, (v) possibility to store the performed jobs, and (vi) automated export of the results in various formats. PREDetector 2.0 is available at http://predetector.fsc.ulg.ac.be/.

## INTRODUCTION

Regulatory DNA-binding proteins control gene expression by interacting with short DNA motifs (*i.e* the *cis* elements or transcription factor-binding sites (TFBS)) which prevents (repressor) or instead assists (activator) RNA polymerase binding to promoter boxes and thus the initiation of transcription. The genome-wide distribution map of the *cis*-acting elements of a TF (or ‘regulon’) provides an estimation of the spectrum – wide or narrow – of its targeted processes. At a broader scale, unveiling the *cis*-*trans* regulatory networks is key for understanding the behaviour, the developmental and physiological capabilities of living organisms. Reduced costs associated with whole genome sequencing (WGS) together with available collections of known TFBSs in specialized databases made the computational prediction of regulons popular prior to any other kind of genome-wide investigation in domains related to Systems Biology (1). One appropriate way to predict the occurrence of *cis* elements belonging to a specific regulator in a given genome is performed via a so-called supervised motif finding approach (2) where a position weight matrix (PWM) is created from aligned known – experimentally validated – binding sites of a TF. The matrix is then used to scan a genome to identify DNA-motifs (also referred to ‘hits’) identical or similar to the training set of sequences used to generate the PWM, and thus potentially bound by the TF of interest.

In 2007 we presented the PREDetector (Prokaryotic Regulatory Elements Detector) software (3) which at that time was, and still is, the first tool that simultaneously provided a threshold-free motif finding approach combined with a statistical means to estimate the reliability of the predicted hits. In a recent opinion article (4), we discussed the necessity to propose web tools in which users can freely adjust the screening parameters and exposed the conflicting objectives of software authors (bioinformaticians) and software users (biologists). Indeed, developers aim at providing conservative tools with reliability criteria as their main priority (bias towards specificity) and thus propose programs limiting the output to the very best matches (high threshold scores or low P-values), while biologists aim to plan investigations based on their own call and are worried of missed opportunities (bias towards sensitivity). We illustrated how disastrous such “absolute truth above, absolute falsehood below” practice may sometimes be, with a series of *cis*-*trans* relationships that could only be predicted – and later experimentally validated – by tools that do not impose stringent screening parameters (4).

Since we strongly believe that regulon prediction cannot rely only on fully automated and statistically thresholded analyses, our aim was to develop an interactive tool inheriting the philosophy of PREDetector, but allowing the user to adjust each of the relevant parameters of the regulon prediction and immediately visualize the results in a convenient and dynamic way. Because of this need for a fully interactive tool and the highly (daily) evolutive nature of the underlying data used by PREDetector (genome sequences and repositories that host them), we decided to focus on the development of an online tool, PREDetector 2.0, presented in this paper.

## IMPLEMENTATION

PREDetector 2.0 was mostly written in R (5) with server-side TFBS prediction modules making use of BioStrings (6) and IRanges (7) packages. The creation of PWM logo uses the Seqlogo package (8). The client-side interface was developed as a web-based application built on top of the Symfony2 PHP framework (<u>http://symfony.com</u>). Interactive features are implemented with JavaScript libraries, including jQuery (<u>http://jquery.com</u>) and d3.js (<u>http://d3js.org</u>). The underlying database is built and managed with SQLite3 (<u>http://www.sqlite.org</u>).

## RESULTS

### Genome and matrix selection/creation, and screening parameters

The online PREDetector interface is freely accessible but requires a preliminary registration because a workspace has to be created for each individual user where they can store genomes, PWM and prediction results. The main panel of the webtool is divided into three tabs, namely Sequences, Matrices, and Regulon predictions, which can be seen as the PREDetector 3-step workflow. Video tutorials to assist users in learning how to use the software, to edit the screening parameters, interpret and to save the prediction results are available directly from the ‘Help’ section of the software web interface.

From the first tab – Sequences (Figure 1) – users can view, select, and download bacterial genomes from the daily updated repository databases. The user can choose to use genome sequences and annotations available in two public repositories: the GenBank repository of the National Center for Biotechnology Information (NCBI) (9), and the Pathosystems Resource Integration Center (PATRIC) (10). Since the regulon prediction quality assessment relies on the identification of the genomic context of identified TFBS (coding vs non-coding regions, see below), only genomes featuring valid annotations can be downloaded. Genomes that successfully passed this quality check are downloaded and stored for further analyses in the user's private collection called ‘My Sequences’.

**Figure 1.**
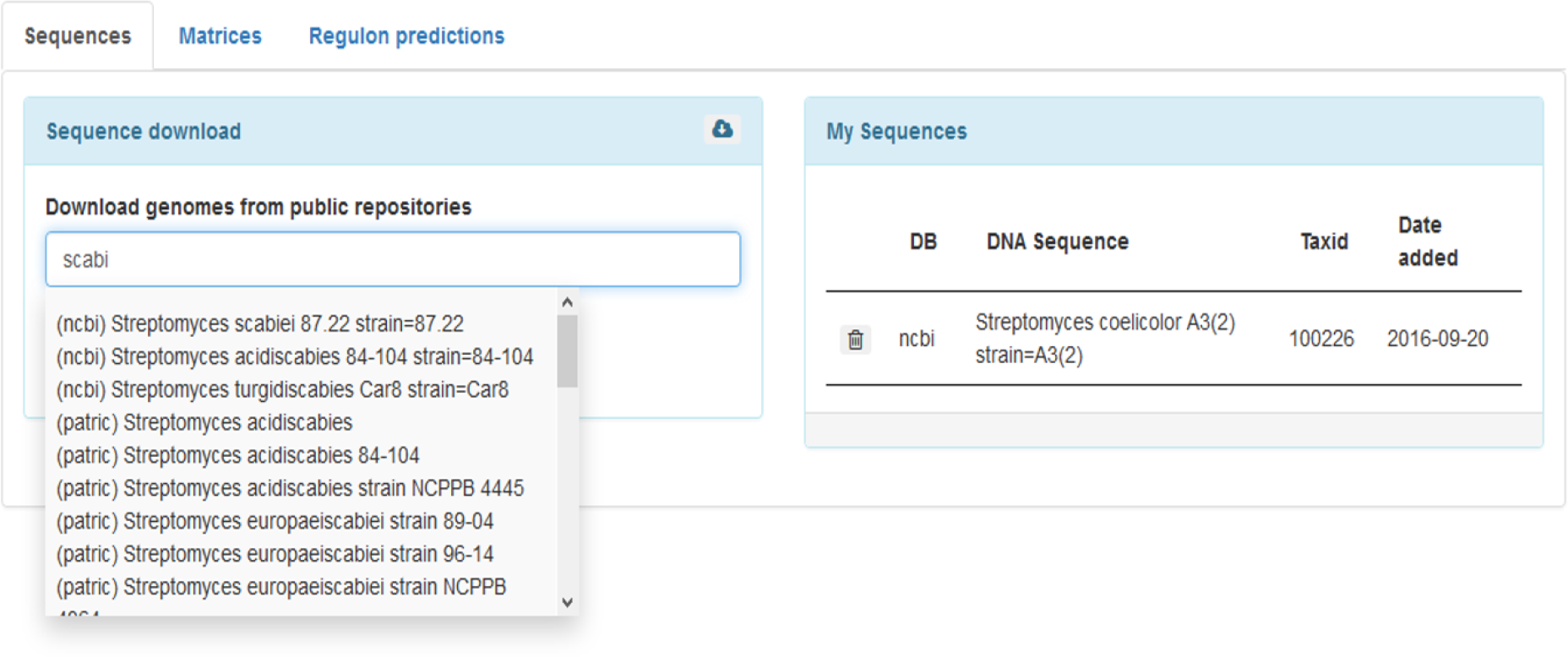
The *Sequences* tab. Users can retrieve genome sequences and information from NCBI and PATRIC databases and are allowed to store genome sequences in their private profile (*My Sequences*) for further analyses.

In the second tab called Matrices (Figure 2), users can either select a list of sequences associated with their TF of interest that are available from public repositories (11), or create their own PWM by providing a set of same-length sequences, in FASTA format. The downloaded or created PWMs can be stored in the user's private collection called ‘My Matrices’ for subsequent use, and a weblogo associated with each saved matrix is generated using the SeqLogo package (8) (Figure 2).

**Figure 2.**
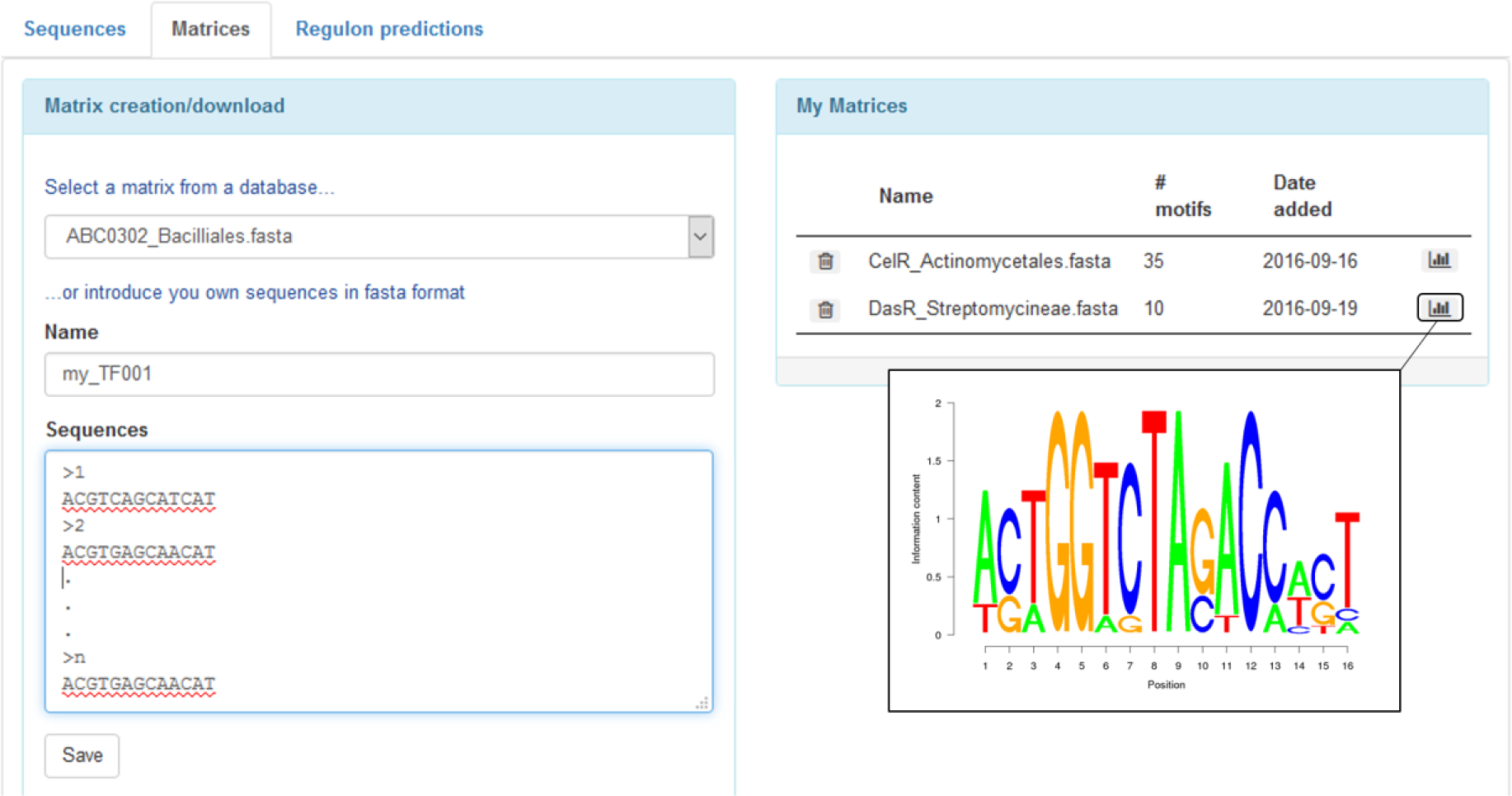
The *Matrices* tab where to download or create a PWM. Users that choose to create their own matrix have to provide a name for the PWM and enter a minimum of two sequences of equal length in FASTA format. Created or downloaded matrices appear in the box on the right called ‘*My Matrices*’ and become available for further analyses. Highlighted, the icon to visualize the weblogo associated with the training set of sequences used to generate the PWM.

At the Regulon predictions tab (Figure 3), two main screening parameters can be specified prior to starting the DNA-motif identification work i.e., the distance range for co-transcription between adjacent genes (operon prediction), and the distance range upstream the translational start codon at which users expect to find functional *cis*-acting elements (Figure 3). Once these parameters are set, users select a genome/matrix couple and ask the software to start identifying DNA motifs in the organism of interest. Scores calculated for the putative DNA-motifs discovered by the analysis are based on PWM generated according to the formula previously described in PREDetector 1.0 (3) as detailed by Hertz and Stormo (12). When the job is performed, the raw data are stored in the box called ‘My predictions’ (Figure 3).

**Figure 3.**
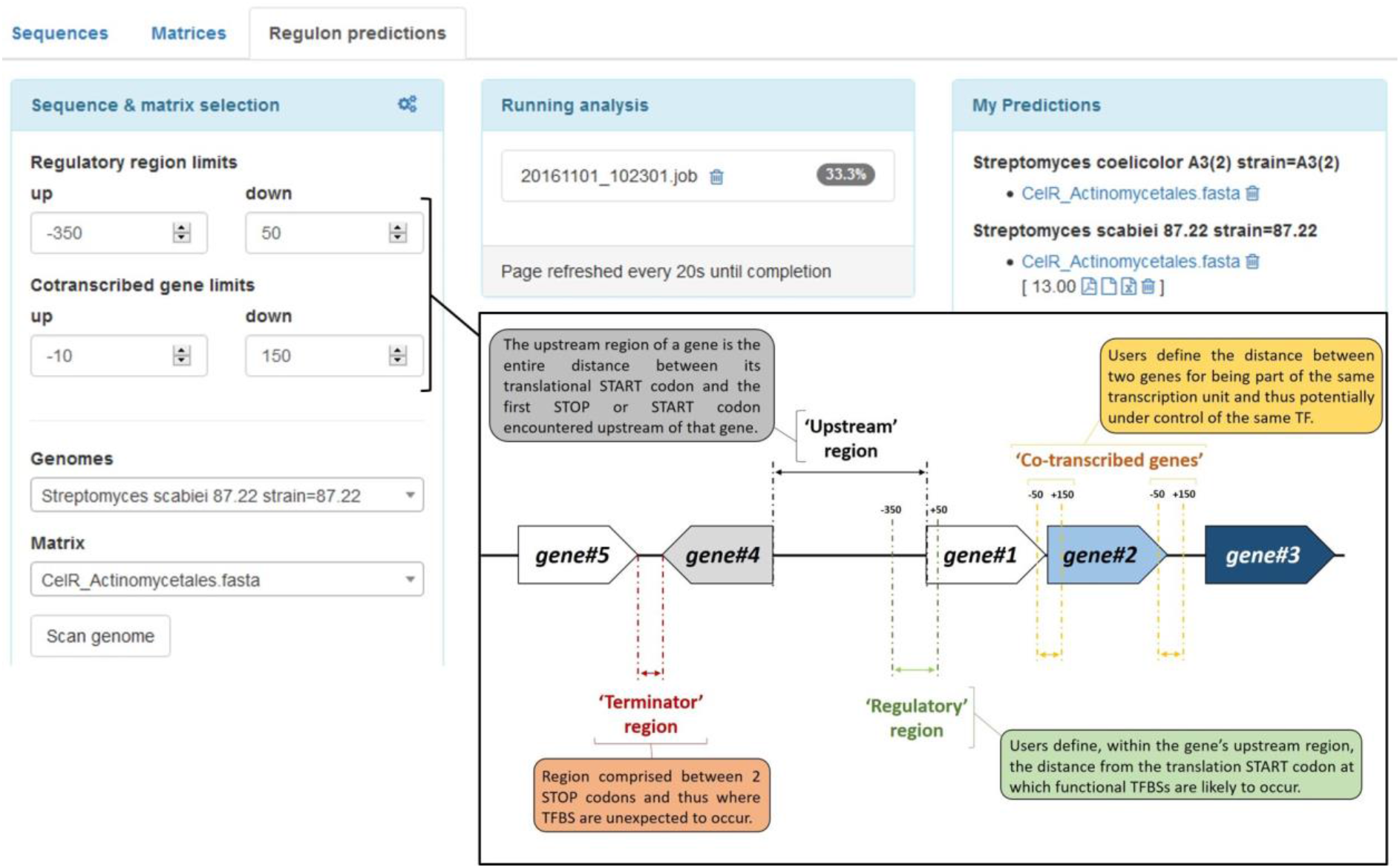
The *Regulon predictions* tab. On the left box, the screening parameters to specify (i) the distance range upstream the translational start codon at which users expect to find functional *cis*-acting elements (Regulatory region limits), and (ii) the distance range expected for co-transcription between adjacent genes (Cotranscribed gene limits) (see right panel for definition of the different ‘regions’). *Running analysis* box = job in progress. *My predictions* box = stored raw data associated with one genome/matrix couple.

### Data visualization and options related to the output list

Clicking on a job stored in ‘My predictions’ box allows users to access the prediction results. The new version of PREDetector proposes two interactive graphical outputs (Figure 4), on top of the table that lists the identified hits, which are by default ranked according to their descending score (Figure 5).

**Figure 4.**
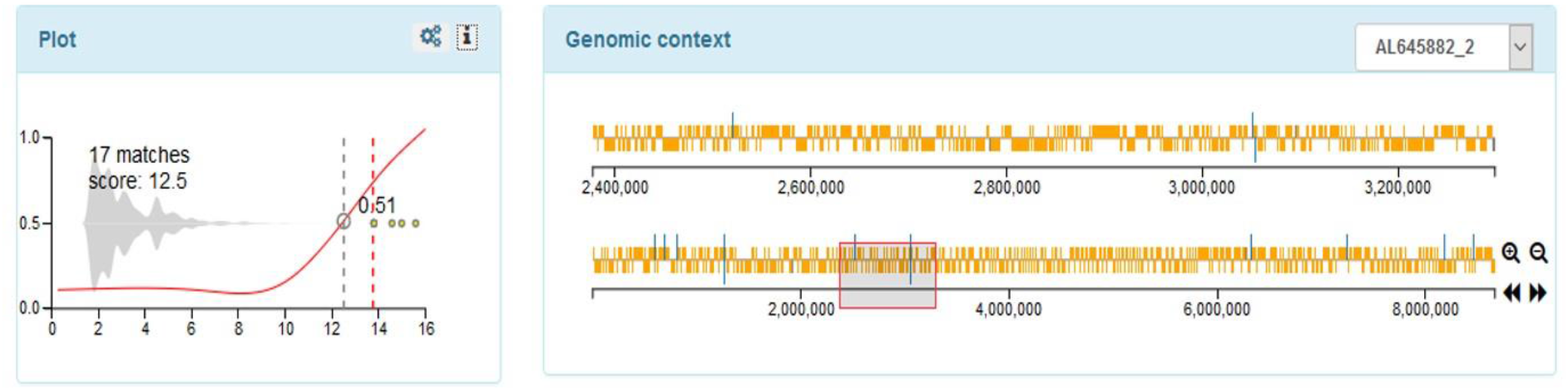
Interactive output graphs of the regulon prediction results. Plot, left panel. Grey bean plot = total number of hits according to the score. Yellow circles = positions of sequences used to generate the PWM. Red line (*reliability curve*) = % of hits in a certain region (intergenic versus coding regions) according to the score. The red dotted line = ‘T_MS_’, for threshold set by default based on the minimal score of the sequences used to generate the PWM. Genomic context, right panel. In orange the coding sequences and in blue the predicted TFBS. The bottom graph represents the full genome sequence and can be used for zooming in a specific region which is then displayed in the top graph.

**Figure 5.**
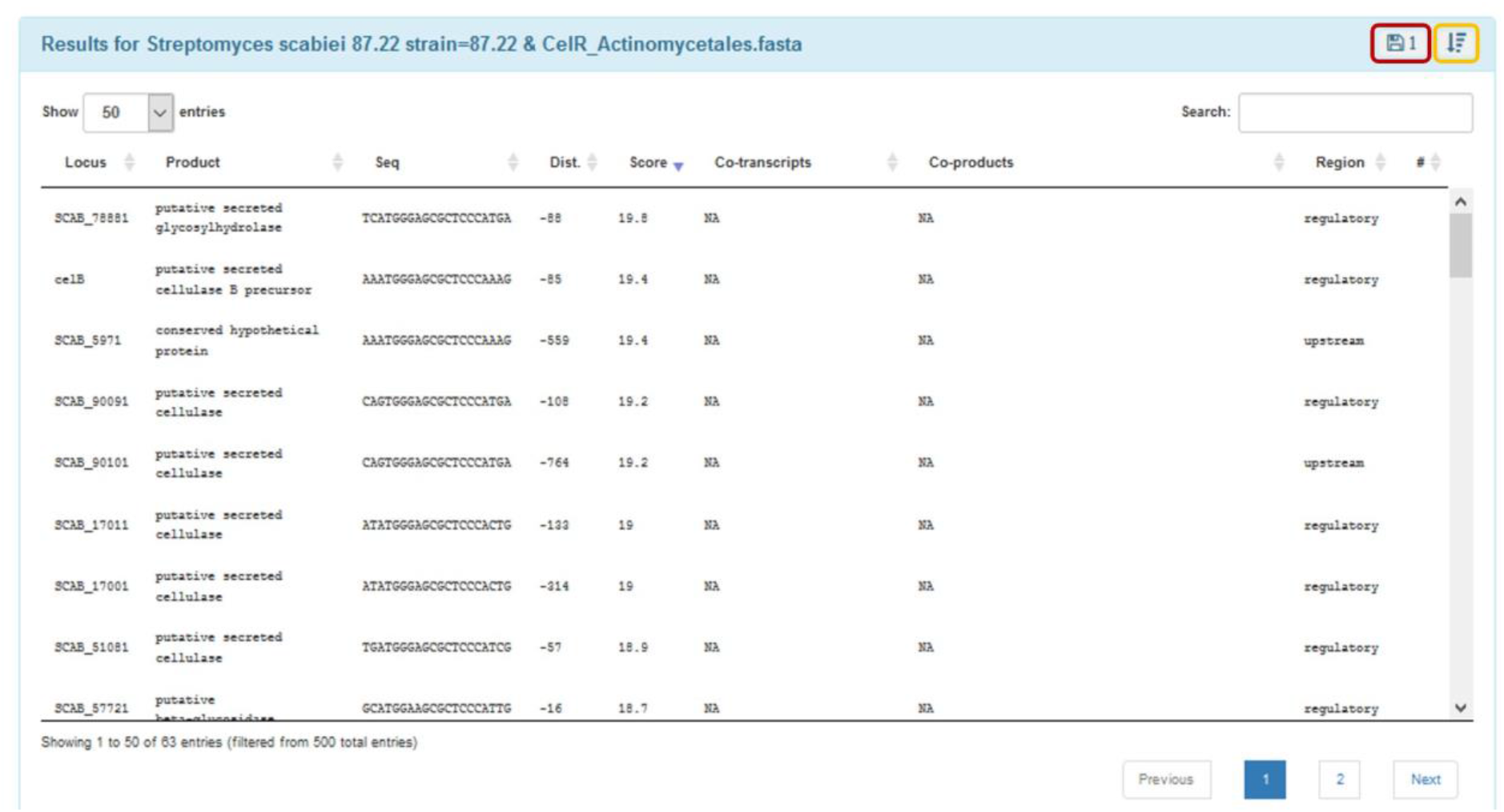
Table displaying the output list of the regulon prediction results. The content of this list is updated dynamically with any changes the users make in the two other interactive boxes of the window (see above Plot and Genomic context graphs). This table is also interactive since users can order the results by clicking on up/down arrows in any column, search for text (top right search field) or click on any row to update the genomic context and zoom on the selected TFBS location. The Floppy disc icon allows users to export a detailed report of their work in various formats (Saved results are stored in ‘My Predictions’ box, see Figure 3). The ‘downwards arrow’ icon allows to display the output list by grouping hits that belong to the same intergenic region (see supplementary Figure S2).

The first panel (Figure 4, left panel called ‘Plot’) displays a bean plot graphic showing the number of hits at each score. The red line in the bean plot – the reliability curve – displays the proportion of hits that localize in the intergenic versus the coding regions, at each score. This line should be considered as a way for users to estimate the probability of a predicted hit to represent a truly functional TFBS. Indeed, since the localization of functional *cis*-acting elements is biased towards intergenic regions and, more precisely, towards the (noncoding) upstream regions, predicted TFBS found at scores at which most of hits are found within coding sequences are likely to include a large number of false positives. PREDetector automatically calculates three thresholds corresponding to specific points of this ‘reliability curve’: ‘T_50_’, at which half of the hits are located in intergenic regions, ‘T_RC_’, which corresponds to the inflection point of the curve below which false positives hits are expected to start to accumulate much faster to the expense of true positives hits, and ‘T_RL_’, which is located at the point where the curve first reaches the minimum imposed by the coding vs non-coding region ratio of the analyzed genome (13). By default, the cut-off score is fixed to a fourth pre-calculated threshold, called ‘T_MS_’, which represents the minimal score of experimentally validated sequences used to generate the PWM. The significance of the different types of score thresholds is described in Supplementary Figure S1.

The bean plot is interactive: users can adjust the threshold as they wish, with immediate adaptation in the Genomic context graphs (see below) and in the output list presented as a table that ranks the predicted hits according to their score (Figure 5). The currently active cut-off score is represented as a vertical dashed red line while previously used (and saved, see below) thresholds for the same genome/PWM couple are represented as dashed green lines. The second panel displays the localization of the hits on the target genome (Figure 4, right panel ‘Genomic context’). Two lines representing the genome are drawn. The open reading frames and TFBSs are displayed as yellow and blue boxes, respectively. The position of the boxes relative to the line representing the chromosome is indicative of the strand where they are located: above the line for positive strand and below for negative strand. The bottom line always displays the full genome sequence with coding sequences and the position of the identified hits. This part of the graph is interactive, and users can select a zone of the genome to be zoomed in, which will be displayed in the upper graph. As for the bean plot, any action on this genomic context plot will immediately be echoed in the result table: only TFBSs located in the defined region will be shown. Conversely, by selecting any row of the result table, the genomic context plot will be updated to focus on the 10,000 bp region flanking the selected TBFS.

### PREDetector result storage and reporting

PREDetector 2.0 offers users the possibility to store their analyses and access their reports in various formats (see ‘My predictions’ box in Figure 3). By clicking on the floppy disk button in the header bar of the result table panel (Figure 5), the prediction results are saved as a filtered list of TFBSs whose scores are greater or equal to the active cut-off score, as represented by the vertical dashed red line in the bean plot. These saved results are labelled with their threshold score and accessible from the ‘My Prediction’ panel in the ‘Regulon Prediction’ tab (Figure 3). The raw tabulated data are available as a tabulated text file (.tsv) or a Microsoft Excel file (.xlsx), and a more complete analysis report is produced as a PDF file. The first part of the PDF report summarizes the input information and the selected screening parameters (PWM, cut-off score, weblogo, genomic distribution, bacterial species…), while the second part of the report displays the table that lists the putative binding-sites and the associated genes predicted to be bound/controlled by the TF under study (supplementary data S3).

## ACKNOWLEDGEMENTS

MH and SR are research associates at the Belgian Fund for Scientific Research (F.R.S-FNRS). All authors are thankful to users of version 1.0 who expressed their wishes to see PREDetector to live on and their advices on how to improve it. We are also grateful to Pierre Martin for his assistance in editing the video tutorials for the PREDetector Help (https://www.youtube.com/channel/UC8VmzdJXMVTzXPb6ryeBY7w).

## Supplementary data

**Supplementary Figure S1.**
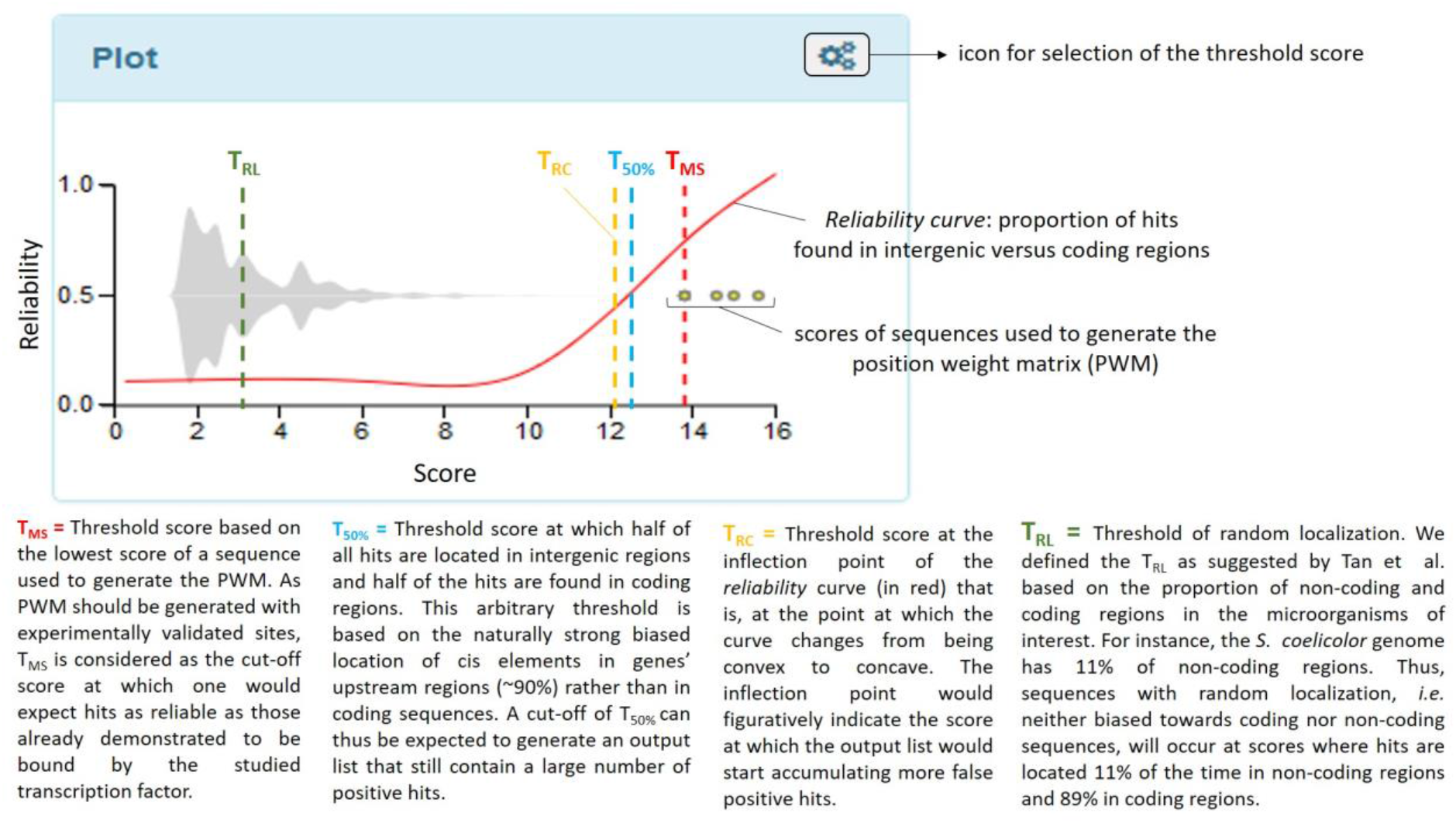
Definition of the different types of score thresholds.

**Supplementary Figure S2.**
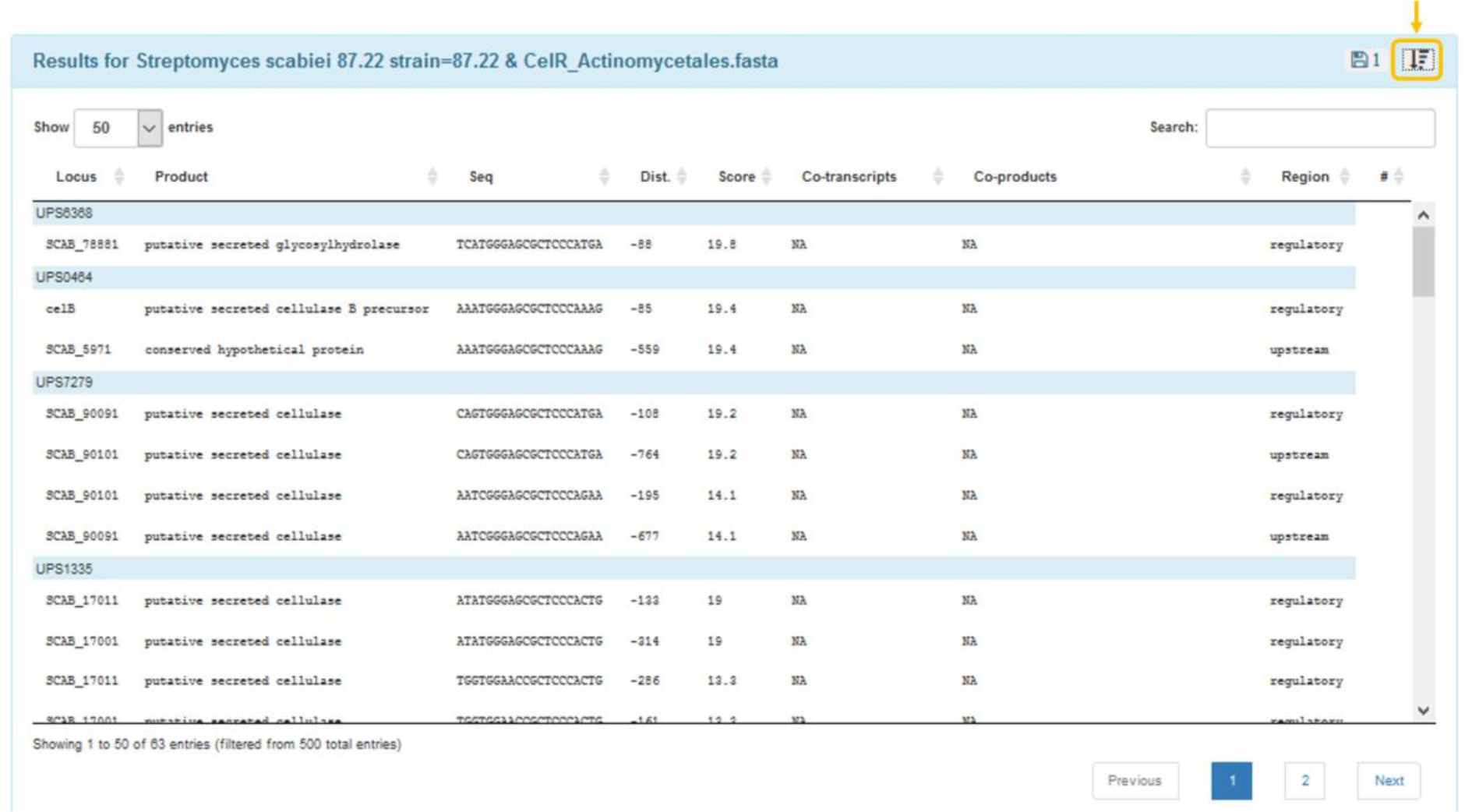
Table with predicted hits identified in the same upstream region grouped together. Clicking on the highlighted icon will rearrange hits according to their score and their localization.

**Supplementary data S3.**

Example of Regulon Prediction report provided as separate file from the manuscript.

